# IP_3_R activity increases frequency of RyR-mediated sparks by elevating dyadic Ca^2+^

**DOI:** 10.1101/2020.08.13.249144

**Authors:** Joshua Chung, Agnė Tilūnaitė, David Ladd, Hilary Hunt, Christian Soeller, Edmund J. Crampin, Stuart Johnston, H. Llewelyn Roderick, Vijay Rajagopal

## Abstract

Calcium (Ca^2+^) plays a critical role in the excitation contraction coupling (ECC) process that governs the contraction of cardiomyocytes during each heartbeat. While ryanodine receptors (RyRs) are the primary Ca^2+^ channels responsible for mediating cell-wide Ca^2+^ transients during ECC, Ca^2+^ release via inositol 1,4,5-trisphosphate (IP_3_) receptors (IP_3_Rs) have been reported to elicit ECC-modulating effects. Recent studies suggest that the proximal localization of IP_3_Rs at dyads grants their ability to modify the occurrence of Ca^2+^ sparks (elementary Ca^2+^ release events that constitute ECC-associated Ca^2+^ transients) which may underlie the modulatory effects on ECC. Here, we aim to uncover the mechanism by which IP_3_Rs affect Ca^2+^ spark dynamics. To this end, we developed a mathematical model of the dyad that incorporates IP_3_Rs to reveal their impact on local Ca^2+^ handling and corresponding Ca^2+^ spark formation. Consistent with published experimental data, our model predicts that the propensity for Ca^2+^ spark formation increases with IP_3_R activity. Our simulations support the hypothesis that IP_3_R activity elevates Ca^2+^ within the dyad, sensitizing proximal RyRs for future release. However, this lowers Ca^2+^ in the JSR available for release and thus results in Ca^2+^ sparks with the same duration but lower amplitudes.

## 1 Introduction

Underpinning the heart’s pumping action is the concerted contraction and relaxation of individual cardiomyocytes, governed by the excitation-contraction coupling (ECC) process. In ventricular cardiomyocytes, ECC is initiated at the arrival of an action potential (AP) that permits a small calcium (Ca^2+^) influx through voltage-gated L-type Ca^2+^ channels (LTCCs) into 10 – 15 nm wide microdomains delimited by T-tubules and the junctional cisternae of the sarcoplasmic reticulum (SR) (**Figure 1**). The Ca^2+^ influx into these microdomains (henceforth dyads) induces a larger Ca^2+^ release from the SR via resident ryanodine receptors (RyRs). This Ca^2+^-induced Ca^2+^ release (CICR) raises local Ca^2+^ concentration ([Ca^2+^]) in the dyad, producing elementary Ca^2+^ release events that underlie ECC known as Ca^2+^ sparks (1,2). By virtue of the distribution of T-tubules at 1.8 μm intervals across the cardiomyocyte that penetrate throughout the cell volume forming dyads, the synchronous evocation of Ca^2+^ sparks at dyads by an AP facilitates the transient global rise in cytosolic Ca^2+^ levels. This allows sufficient Ca^2+^ to bind to troponin C (TnC) in myofilaments, thereby enabling the cross-bridge cycle that contracts the cardiomyocyte (3).

**Figure 1.**
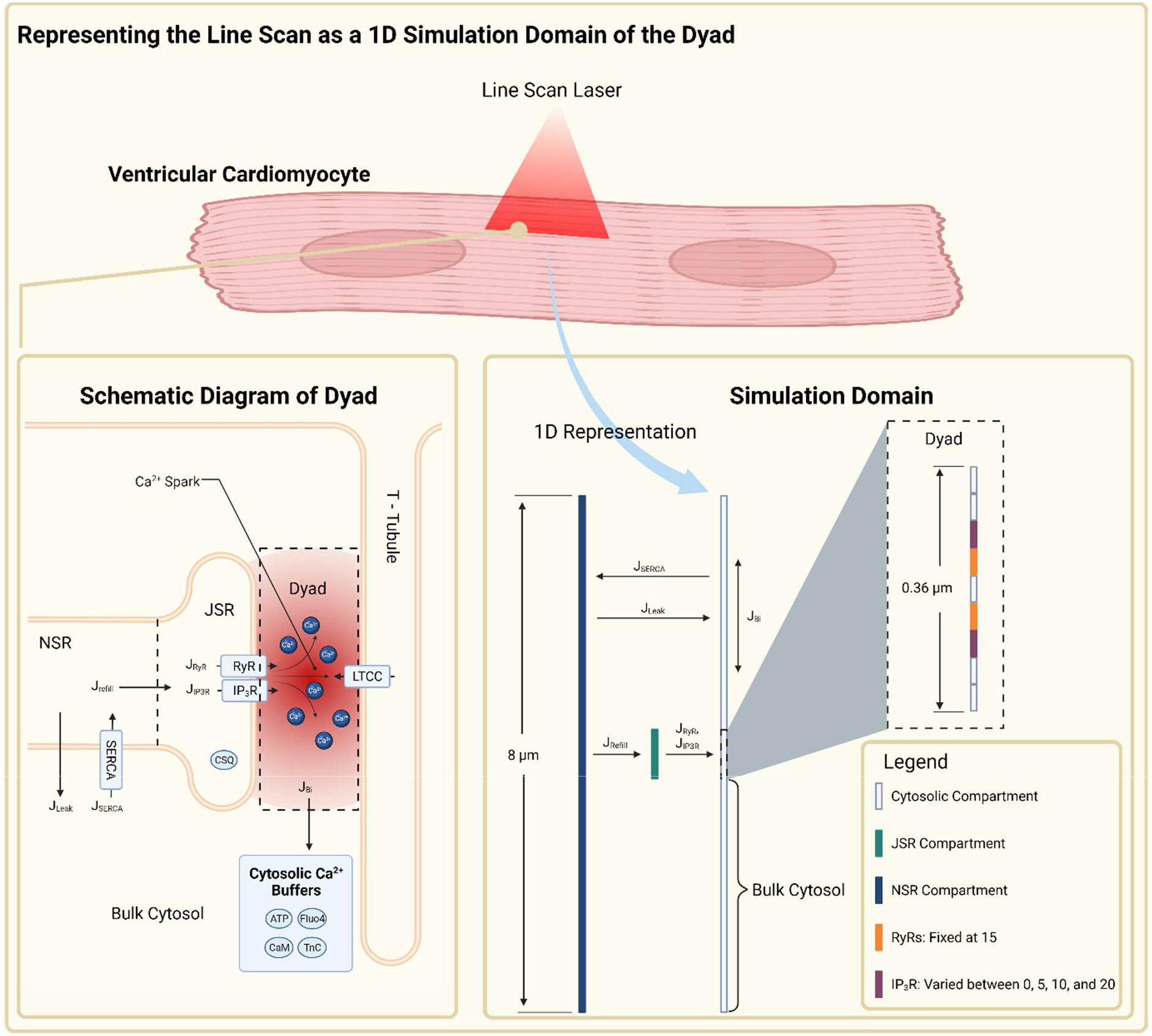
Schematic diagram of the compartments, fluxes, and arrangement of Ca^2+^-handling proteins in the dyad and its 1D representation in the model. The dyad is represented on an 8-μm, 1D computational domain with three compartments: cytosol, JSR and NSR. The center nine elements of the cytosolic compartment represent the dyadic region where RyRs and IP_3_Rs are located while the remaining elements represent the bulk cytosol where Ca^2+^ is additionally subject to J_SERCA_, J_Leak_ and J_B_4__. Ca^2+^ from the JSR is released in the dyadic region via open RyRs and IP_3_Rs and diffuses along the cytosolic compartment, reacting with buffers before eventually being pumped back into the NSR which refills the JSR.

Like RyRs, inositol 1,4,5-trisphosphate (IP_3_) receptors (IP_3_Rs) are another family of Ca^2+^-regulated Ca^2+^ channels that reside on the SR of cardiomyocytes (4). IP_3_Rs are positively regulated by their ligand, IP_3_ (5), which is produced following phospholipase C activation and phosphatidylinositol 4,5-bisphosphate hydrolysis downstream of G protein-coupled receptors (GPCRs) as well as certain receptor growth factor receptors (6). Indeed, ventricular cardiomyocytes stimulated by G_q_ coupled GPCR agonists, such as endothelin-1 (ET-1), lead to IP_3_-induced Ca^2+^ release (IICR) via IP_3_Rs, which are shown to promote ECC-modulating effects such as arrhythmia and positive inotropy (7–12).

Despite lower expression levels (13) and Ca^2+^ conductance (5) relative to RyRs, IP_3_Rs elicit these ECC-modulating effects by their localisation to functionally relevant Ca^2+^ signalling sites in the cell (14). A notable example is the colocalization of IP_3_Rs and RyRs at dyads (8,15). It has been recently shown that stimulating the activity of IP_3_Rs significantly increases the frequency of dyadic Ca^2+^ spark events (15). In this regard, IICR is hypothesised to elevate dyadic Ca^2+^, thereby priming and recruiting otherwise “silent” RyRs towards activation (8,11,14,15). The resulting increase in Ca^2+^ spark propensity may then lead to ECC-modulating effects observed (11,16).

Here, we employed computational modelling to simulate the effects of IICR in the dyad. We developed a 1D spatial model of a dyad containing RyRs and type 2 IP_3_Rs. Using this model, we varied the number of IP_3_Rs in the dyad and simulated its effect on the local Ca^2+^ dynamics as well as the properties of Ca^2+^ Ca^2+^ sparks generated. Our model predicts that increasing IP_3_R activity increases the baseline dyadic [Ca^2+^] at the expense of that in the JSR. This elevation of dyadic [Ca^2+^] then sensitises colocalized RyRs to be more active, consequently increasing the propensity of Ca^2+^ spark formation. The decrease in JSR [Ca^2+^] thus resulted in Ca^2+^ sparks with lower amplitudes but a similar duration.

## 2 Methods

### 2.1 Model Formulation

We model the spatiotemporal evolution of Ca^2+^ as a system of partial differential equations (PDEs) at three interconnected compartments: cytosol, junctional SR (JSR), and network SR (NSR). The spatiotemporal evolution of [Ca^2+^] in these compartments is described by the variables [*Ca*^2+^]*_c_*, [*Ca*^2+^]*_JSR_*, and [*Ca*^2+^]*_NSR_* respectively. These are shown in order in the equations below:

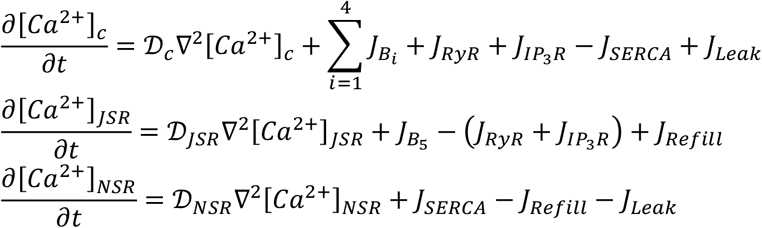

where 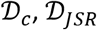, and 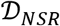 represent the diffusivity of Ca^2+^ in the cytosol, JSR, and NSR compartments, respectively. *J_RyR_* and *J*_*IP*_3_*R*_ correspond to the Ca^2+^ release fluxes by open RyRs and IP_3_Rs respectively. *J_SERCA_* corresponds to the Ca^2+^ uptake flux by sarco/endoplasmic reticulum Ca^2+^ ATPase (SERCA). *J_Refill_* corresponds to the Ca^2+^ refill flux from the NSR into the JSR compartment. *J_B_i__* corresponds to the flux of Ca^2+^ binding to mobile and immobile buffer species *i*.

The reaction diffusion of Ca^2+^ buffers are described by

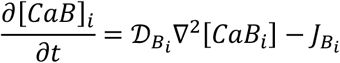

where [*CaB*]*_i_* corresponds to the concentration of Ca^2+^-bound buffer species *i*, with *i* ∈ {1,2,3,4,5} representing buffers ATP, calmodulin (CaM), Fluo-4, troponin C (TnC), and calsequestrin (CSQ) respectively. 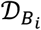 corresponds to the diffusivity of Ca^2+^-bound buffer species *i*. Immobile buffers TnC and CSQ have 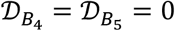.

### 2.2 Calcium Fluxes

The flux for each buffer species *i* is given by

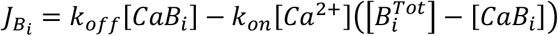

where 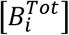 corresponds to the total concentration of buffer species *i*. *k_on_* and *k_off_* corresponds to the forward and backward reaction rates of buffer species *i* with Ca^2+^ respectively.

The refill flux from the NSR to the JSR compartment is given by

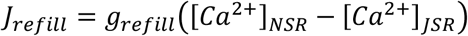

where *g_refill_* is the refill flux rate adjusted to achieve a realistic JSR refill time constant of ~130 ms (17–19) in simulations where the number of IP_3_Rs is 10 (see **Figure 1**). We assume the average number of IP_3_Rs in a cluster to be 10.

The release fluxes from RyRs and IP_3_Rs are given by

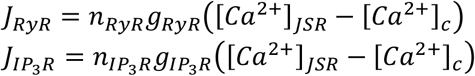

where *n_RyR_* and *n*_*IP*_3_*R*_ correspond to the number of open RyRs and IP_3_Rs, respectively, whereas *g_RyR_* and *g*_*IP*_3_*R*_ correspond to the flux rate of RyR and IP_3_R release, respectively. The value of *g_RyR_* was adjusted to yield a characteristic Ca^2+^ spark profile in the simulation condition where only RyRs are present in the dyad. *g*_*IP*_3_*R*_ is less than *g_RyR_* by a factor of 2.85 as the Ca^2+^ conductance of IP_3_Rs is estimated to be lower than that of RyRs by that amount (5).

Fluxes due to SERCA uptake activity were directly adapted from (20), which takes the form

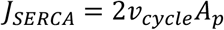

where *v_cycle_* is the cycling rate per SERCA molecule and *A_p_* is the cytosolic concentration of SERCA homogenously spread throughout the bulk cytosolic region. The complete expression of each term is provided in Supplementary Materials.

An SR leak flux was also introduced to maintain the cytosolic Ca^2+^ background concentration of 0.1 μM. We use the same formulation as the SERCA model to balance *J_SERCA_* such that [*Ca*^2+^]*_c_* does not fall below 0.1 μM. Therefore, the SR leak flux is expressed as,

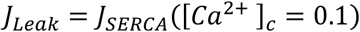

All parameter values are listed in Supplementary Materials.

### 2.3 Calcium Channels

Two types of Ca^2+^ channels are stochastically simulated in the model – RyRs and IP_3_Rs. The gating of each RyR is directly adapted from the 2-state RyR model developed in (21); IP_3_Rs are modelled after the 6-state Siekmann model that incorporates non-steady state kinetics developed and used in (22,23). Mathematical expressions of the IP_3_R model used in (22,23) were parameterised specifically to fit the open probability curve of type 1 IP_3_Rs (IP_3_R-1). IP_3_R-1 have different channel activities for the same range of [Ca^2+^] compared to type 2 IP_3_Rs (IP_3_R-2) (compare **Figure 8**C and **Figure 8**D in Supplementary Materials), the isoform most expressed in cardiomyocytes (5). Therefore, we varied coefficients and exponents of the IP_3_R model in (22,23) to fit the open probability curve of IP_3_R-2 data from (24). This allowed us to obtain an IP_3_R model specific to IP_3_R-2. This modification is essential due to the higher levels of Ca^2+^ required for channel inhibition, which thus allow IP_3_R-2 to remain active for longer in the conditions of the dyad. Details are described in Supplementary Materials. In simulations containing IP_3_R-2s, their gating behaviour were computed with a fixed IP_3_ concentration of 0.15 μM, similar to that used in (22).

### 2.4 Model Geometry

The dyad and its surrounding cytoplasmic space are represented on a 1-dimensional (1D) simulation domain of size 8 μm. The 1D simulation domain reflects the portion of a typical experimental confocal line scan where a dyad is located. The buffering of Ca^2+^ by mobile buffers ATP, CaM, and Fluo-4 take place throughout this domain. The domain consists of 200 elements of size 0.04 μm, with the center nine elements (0.36 μm long) representing the dyadic region where RyRs and IP_3_Rs are placed as shown in **Figure 1**. Elements outside this region represent the bulk cytosol where Ca^2+^ is subject to additional buffering by TnC and uptake into the NSR by SERCA. In all simulations, the number of RyRs in each element is fixed at 15, consistent with the average number of RyRs in a cluster as determined by super resolution microscopy techniques in healthy cardiomyocytes (25–27). However, similar data on IP_3_R clusters are not yet available. Therefore, to investigate the effect of enhanced IP_3_R activity, the number of colocalized IP_3_Rs in their element is varied between 0, 5, 10, and 20, corresponding to situations where there are no, low, intermediate, and high levels of IP_3_R activity relative to the number of RyRs. The JSR compartment is designated the same location and the same number of elements as the dyadic region. Open RyRs and IP_3_Rs thus result in a Ca^2+^ flux from the JSR into the dyad that is driven by the difference in [Ca^2+^] between the two compartments. Ca^2+^ in the JSR is subject to buffering by CSQ and refill from the NSR compartment. The non-junctional regions of the NSR compartment are homogenously distributed with SERCA that pumps Ca^2+^ from the bulk cytosol into the SR. SR leak fluxes are likewise present along non-junctional regions of the NSR compartment and leaks Ca^2+^ into the bulk cytosolic region to maintain a baseline cytosolic [Ca^2+^] of 0.1 μM. No-flux conditions were imposed on the boundaries of the simulation domain.

### 2.5 Numerical Methods and Implementation

The system of PDEs were discretised using the forward time centered space finite difference scheme, similar to (28). The resulting system of ODEs was solved using the explicit Euler method with adaptive time stepping capped at a maximum of 1 × 10^−4^ ms and a regular spatial resolution of 0.04 μm. Stochastic IP_3_R and RyR gating states were solved using a hybrid Gillespie method as described in (29). The time at which any one receptor changes state determines the time step forward for which the system is solved (adaptive time stepping). Each simulation was repeated 200 times. In all simulations, the model was run for 1000 ms to ensure the system achieves steady state before any results were recorded. All codes and computations were implemented in MATLAB (The MathWorks Inc., Natick, Massachusetts).

## 3 Results

### 3.1 1D model reproduces calcium spark dynamics

The first column of **Figure 2**A illustrates the typical time evolution of Ca^2+^ in different compartments of the model during a Ca^2+^ spark in RyR-only simulations i.e., no IP_3_Rs. To replicate CICR during ECC arising following Ca^2+^ influx via LTCCs, Ca^2+^ sparks were initiated by introducing a 2-ms Ca^2+^ flux, reaching ≈ 30 μM, to elements in the dyadic region where RyRs are placed at the 1000 ms time point. This influx can be observed by the initial rise in [Ca^2+^] (with no variance) at the center of the dyadic region. The resultant opening of RyRs occurs rapidly and releases a greater amount of Ca^2+^ from the JSR, thus providing a temporary positive feedback mechanism for the opening of other RyRs via CICR. RyRs open shortly after the initiation trigger and peaked at ≈ 13 RyRs for ≈ 9 ms before closing completely after ≈ 16 ms, consistent with simulation results from (21) whereby RyRs terminate after ≈ 20 ms of activity. During this time, dyadic [Ca^2+^] increased to ≈ 300 μM on average and declined back to ≈ 0.1 μM due to diffusion and chelation by buffers in the cytosol. While in the JSR, [Ca^2+^] declines and reaches its nadir at ≈ 300 μM ≈ 13 ms after the initiation trigger, during which point RyRs have already begun closing. These results support the induction decay mechanism of Ca^2+^ spark termination proposed by (30) whereby the decay of the Ca^2+^ flux through RyRs due to JSR depletion eventually impedes the inter-RyR regenerative CICR during a Ca^2+^ spark, resulting in its termination. [Ca^2+^] in the JSR is then gradually replenished by Ca^2+^ in the NSR at a time constant of ≈ 130 ms in the 10 IP_3_R simulation, consistent with experimental data (18,19). Together, our model depicting the 1D representation of the dyad is thus capable of reproducing Ca^2+^ spark dynamics in reasonable agreement to that reported in other modelling and experimental studies (17–19,21).

**Figure 2.**
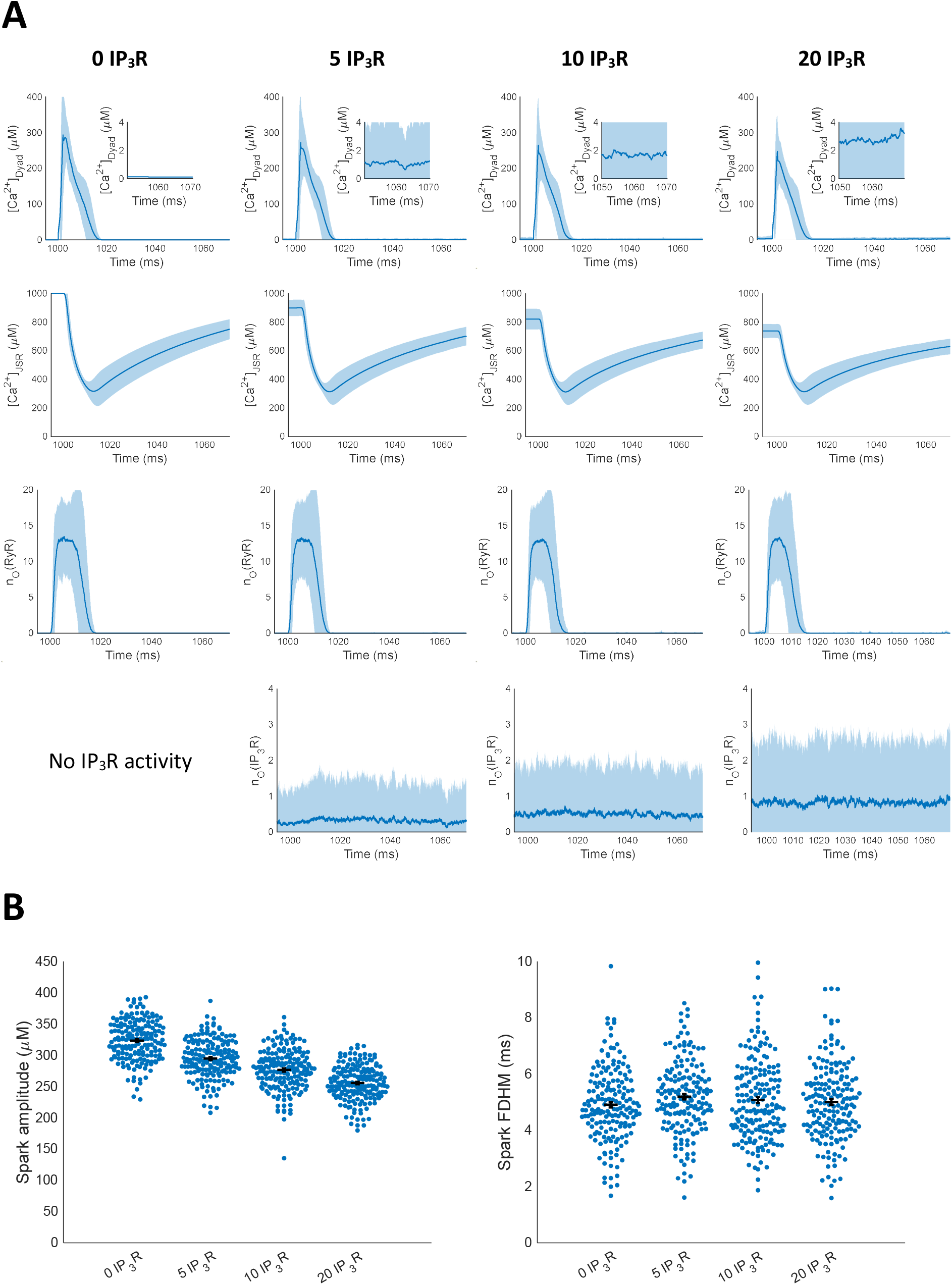
Ca^2+^ dynamics associated with LTCC-initiated Ca^2+^ sparks for different number of colocalized IP_3_Rs. **A:** Time evolution of [Ca^2+^] in the dyad and JSR and the corresponding activity of RyRs and IP_3_Rs during a Ca^2+^ spark. Solid lines and surrounding shaded region show mean and 95% confidence intervals respectively. **B:** Amplitude and FDHM of Ca^2+^ sparks generated based on [Ca^2+^] in the dyad.

### 3.2 Increased IP_3_R-2 activity decreases Ca^2+^ spark amplitude and Ca^2+^ stores

**Figure 2**A illustrates the effect of incorporating an increasing number of IP_3_R-2s at the dyadic region. Despite varying the number of IP_3_Rs in the dyad, our model is able to robustly simulate Ca^2+^ spark events. Due to the significantly faster state transition rates and higher activity at lower [Ca^2+^] of IP_3_R-2s relative to RyRs, incorporating IP_3_Rs in the dyad essentially causes a continuous Ca^2+^ “leak” from the JSR into the dyad. Hence, a 1 s wait time was allocated to allow the system to equilibrate to steady state before simulating any Ca^2+^ release events. This amount of time was sufficient for the system to equilibrate as triggering Ca^2+^ sparks in simulations with longer wait times did not alter the resultant steady state [Ca^2+^]. After equilibration, the average baseline dyadic [Ca^2+^] rose above (inset of first row in **Figure 2**A) while that in the JSR fell below (second row of **Figure 2**A) the model’s initial conditions of 0.1 μM and 1 mM respectively. Moreover, the magnitude of these changes increases with the number of IP_3_Rs present in the dyad. We thus attribute these effects to the increased average number of open IP_3_Rs (fourth row of **Figure 2**A). Altogether, our results suggest that enhancing IP_3_R activity elevates dyadic [Ca^2+^] at the expense of [Ca^2+^] in the JSR.

To test the effect of IP_3_R colocalization on Ca^2+^ spark dynamics, we initiated Ca^2+^ sparks in simulations where IP_3_Rs are present by introducing a Ca^2+^ flux into RyR-containing elements as described earlier. Generated Ca^2+^ sparks have amplitudes that decrease with the increasing number of colocalized IP_3_Rs (**Figure 2**B). This correlates well with the lower JSR [Ca^2+^] available for release at steady state. However, the duration of these Ca^2+^-triggered Ca^2+^ sparks, measured by its full duration at half maximum (FDHM), is not significantly different (**Figure 2**B). This result can also be indirectly inferred from the time to complete closing of RyRs and time to nadir of [Ca^2+^] in the JSR that are not significantly altered with increasing IP_3_R activity. Mechanistically, the elevated dyadic [Ca^2+^] together with the lower JSR [Ca^2+^] at steady state collectively results in RyR Ca^2+^ release fluxes that sustain inter-RyR CICR while depleting the JSR such that the Ca^2+^ spark duration remains unchanged. In all cases, the occurrence of Ca^2+^ sparks coincide with the transient opening of RyRs while the average IP_3_R activity remained relatively constant throughout the simulation. This suggests that RyRs, and not IP_3_Rs, are primarily responsible for the manifestation of Ca^2+^ sparks, which is consistent with experimental results that show an almost complete loss of Ca^2+^ spark events when RyRs are inhibited (15). Our model also successfully reproduced the experimental observation that [Ca^2+^] in the JSR decreases to the same level after a Ca^2+^ spark event regardless of its initial concentration (19), further bolstering our confidence of this model in simulating Ca^2+^ sparks.

### 3.3 IP_3_Rs increase propensity for spontaneous Ca^2+^ sparks

By virtue of elevating dyadic [Ca^2+^], IP_3_Rs may play a role in enhancing the formation of Ca^2+^ sparks. Indeed, cardiomyocytes treated with G_q_ agonists or IP_3_ significantly increased the number of spontaneous Ca^2+^ spark events (7,8,31). This was attributed to IICR, but the underlying mechanism that led to this observation is not fully resolved. To test whether the colocalization of IP_3_Rs in the dyad is a potential cause for the increase in spontaneous Ca^2+^ spark events, we performed simulations in the absence of LTCC initiation – all Ca^2+^ sparks generated occur spontaneously. After a 1 s wait time for system equilibration, the simulation was allowed to run for a further 2 s from which our results were obtained.

We recorded the number of Ca^2+^ spark events generated from these simulations and their associated properties (amplitude and FDHM). We find that the percentage of simulations with at least 1 Ca^2+^ spark event increases with the number of IP_3_Rs (**Figure 3**A). Consistent with triggered Ca^2+^ sparks, the average amplitudes of spontaneously generated Ca^2+^ sparks also decrease (**Figure 3**B) with an increase in the number of colocalized IP_3_Rs while their FDHM remain unchanged (**Figure 3**C). We hypothesise that the increase in spontaneously generated Ca^2+^ sparks is due to the sensitization of RyRs by an elevated dyadic [Ca^2+^]. To verify that RyRs are more active due to their sensitization by IICR, we also recorded the number of RyR openings that did not develop into Ca^2+^ sparks (an example of detecting these events is shown in **Figure 7** of Supplementary Materials). Expectedly, the average number of spontaneous RyR openings that do not lead to the formation of Ca^2+^ spark events also increased with the number of colocalized IP_3_Rs (**Figure 3**D), reinforcing that RyRs are indeed more active in the presence of greater IP_3_R activity. The increased number of spontaneous RyR openings thus raises the probability for Ca^2+^ spark formation and contributes to the decreased JSR [Ca^2+^] at steady state to some degree. Altogether, our results support the notion that IICR via colocalized IP_3_Rs elevates dyadic [Ca^2+^] sensitizing RyRs to promote Ca^2+^ spark formation.

**Figure 3.**
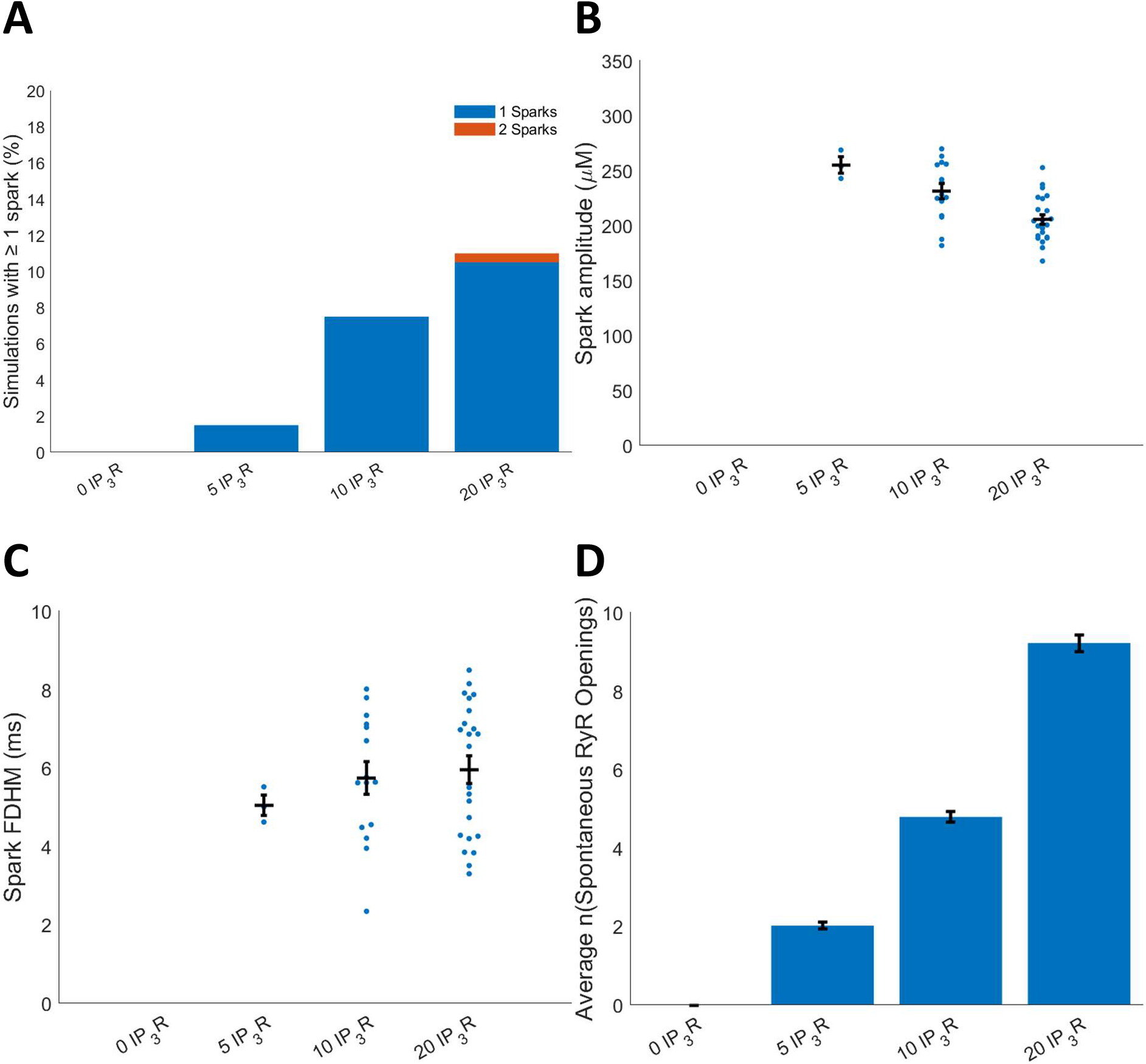
The number of spontaneous Ca^2+^ sparks increase with IP_3_R activity. **A:** Percentage of simulations where more than 1 Ca^2+^ spark event spontaneously occurred and the average number of Ca^2+^ spark events per simulation. **B:** Amplitude of spontaneous Ca^2+^ spark events. **C:** FDHM of spontaneous Ca^2+^ spark events. **D:** Average number of spontaneous RyR opening events that do not lead to a Ca^2+^ spark.

## 4 Discussion

While the increased activity of IP_3_Rs in the cardiomyocyte is shown to elicit ECC-modulating effects (7–12), the mechanistic basis of this action has not been demonstrated. However, evidence increasingly suggests that IICR can modulate ECC by the localisation of IP_3_Rs at functionally important Ca^2+^ signalling sites (14), a quintessential example of which are dyads (8,15). Specifically, the expression of IP_3_Rs at dyads is hypothesised to sensitise native RyRs towards activation via IICR (8,14,15), thus increasing the propensity for the formation of Ca^2+^ sparks – elementary Ca^2+^ release events underlying ECC-associated Ca^2+^ transients. Using a 1D model of the dyad that includes both RyRs and IP_3_R-2s, we set out to test this hypothesis while uncovering the underlying local Ca^2+^ dynamics in the dyad.

### 4.1 IP_3_R-mediated Ca^2+^ release prime RyRs for release

A notable finding in our simulations is that the probability of spontaneous Ca^2+^ spark events increase with the number of colocalized IP_3_Rs. We were also able to uncover the mechanism by which this occurs through our recording of [Ca^2+^] evolution with time at different compartments of the dyad. IP_3_Rs demonstrate vastly different gating behaviour from RyRs. While RyRs are almost always in their closed state at baseline [Ca^2+^], IP_3_Rs are shown to be more highly active, rapidly fluctuating between their open and closed state throughout the simulation time course (fourth row of **Figure 2**A). Consequently, the activity of IP_3_Rs in the dyad is akin to introducing a continuous leak from the JSR into the dyad. Increasing the number of IP_3_Rs increases the magnitude of this “leak”, as can be seen from a lower [Ca^2+^] in the JSR, due to an increased number of open IP_3_Rs on average. This SR [Ca^2+^]-modulating “leak” due to IICR is consistent with that proposed by (32). On the other hand, however, the baseline [Ca^2+^] in the dyad is increased due to IICR (inset in first row of **Figure 2**A). The elevation in dyadic [Ca^2+^], in turn, increases the open probability of colocalized RyRs which is reflected by both the increased number of spontaneous Ca^2+^ sparks (**Figure 3**A) and RyR openings that do not initiate a Ca^2+^ spark (**Figure 3**C). Altogether, our results support the notion that IICR via IP_3_Rs expressed in dyads increases the propensity for RyR-mediated Ca^2+^ spark formation by increasing dyadic [Ca^2+^].

Our findings have important implications about the wider role of IP_3_Rs in cardiomyocytes. As we show that IICR increases the probability of Ca^2+^ spark events by raising dyadic [Ca^2+^], this mechanism may be useful in activating RyR clusters that are usually “silent” during ECC. This can potentially explain the enhanced Ca^2+^ transient amplitude in cardiomyocytes where IP_3_Rs are stimulated (7–11). Indeed, in a recent study in which a dyadic Ca^2+^ reporter was employed IP_3_R activation was found to result in an increase in the number of dyads recruited during ECC (15). In diseased cardiomyocytes, the greater expression of IP_3_Rs (8,12) suggest a compensatory mechanism for the increased decoupling of RyRs from LTCCs due to T-tubule degradation (33,34). But IICR in dyads could also contribute to increased spontaneous Ca^2+^ release events in cardiomyocytes where IP_3_Rs are stimulated, which can have arrhythmogenic consequences (7–11).

### 4.2 Model limitations and implications

We developed a 1D spatial model of a dyad that reproduced all major characteristics of a Ca^2+^ spark. This enabled its utilization in conducting a qualitative investigation into the influence of IP_3_R activity on the dynamics of Ca^2+^ sparks in the dyad. While computationally less expensive, the reduced order of our model from 3D to 1D requires simplifying assumptions that presents several limitations, which we discuss below.

#### 4.2.1 Arbitrary RyR and IP_3_R Placement

In our model, we chose to fix the number of RyRs in a cluster as 15 based on recent estimates obtained from super resolution imaging data (25–27). Since similar data on IP_3_R clusters is unavailable in the literature, the number of IP_3_Rs in a cluster is varied to test the effect of increased IP_3_R activity on the same RyR cluster. These clusters are then arbitrarily placed in elements of the dyadic region as shown in **Figure 1**. The 1D nature of our model precludes our ability to place each RyR in its own element in 3D space such that it can detect Ca^2+^ that has diffused from other RyRs in the cluster. RyRs and IP_3_Rs that belong to the same cluster are placed in one element such that all Ca^2+^ channels in that element are assumed to detect the same dyadic [Ca^2+^]. Similar assumptions have also been employed in previous modelling studies simulating Ca^2+^ sparks (20,35,36). While we acknowledge that developing models of higher dimensions permits one to incorporate the spatial arrangement of individual RyRs in the dyad, which influences Ca^2+^ spark fidelity (17,27), our reduced-order model is sufficient for our purposes of illustrating the general effect of increasing IP_3_R activity on Ca^2+^ spark dynamics and derive an underlying mechanism for its increased occurrence.

#### 4.2.2 Visualisation of Ca^2+^ Spark Fluorescence

Ca^2+^ spark characteristics obtained from experiments are derived from the fluorescence measurement of indicator dyes. To corroborate experimental observations with modelling results, many mathematical models incorporate the reaction kinetics of the indicator dye and simulate the resulting fluorescence measurement. Although we included the reaction kinetics of Ca^2+^ with the indicator dye Fluo-4 in our model, we could not reliably corroborate its fluorescence with experimental observations, which show that Ca^2+^ spark amplitudes do not change in the presence of IICR (15). Contrastingly, our modelling results show progressively decreasing Ca^2+^ spark amplitudes with increasing IP_3_R activity, which we attribute to the decreased JSR [Ca^2+^] available for release.

Whether this decrease in Ca^2+^ spark amplitude, measured in terms of [Ca^2+^], is noticeably reflected in the fluorescence equivalent measurement cannot be directly confirmed from our model. This is because we find that the rise in dyadic [Ca^2+^] during a Ca^2+^ spark saturates the indicator dye, resulting in a plateau of the fluorescence trace (see **Figure 8**A in Supplementary Materials). The saturation of Fluo-4 and the consequent plateau of the fluorescence trace thus prohibits us from conclusively inferring the amplitude and FDHM of the Ca^2+^ spark. We partly attribute this saturation to the diffusion of Ca^2+^ that is restricted to one dimension due to our model design. Additionally, the 1D setup of our model limited our ability to convolve the simulated fluorescence with a point spread function (PSF) of the objective lens of a confocal microscope. Hence the 1D model precludes a realistic visualisation of Ca^2+^ sparks as they would be experimentally observed. We acknowledge this as a major limitation of our model. Nevertheless, simulated spark properties in in our model (**Figure 2**A) is consistent with many previous experimental and model findings. Hence, we base our findings on [Ca^2+^] instead.

Besides our model limitations, we also consider the discrepancy between our modelling results and experimental data to be partially attributable to reaction kinetics between the dye and dyadic Ca^2+^. The dye does not fully provide a measure of dyadic Ca^2+^ due to the dissociation constant between Ca^2+^ and the indicator dye. Therefore, our model results that show decreasing Ca^2+^ spark amplitude with increased IICR activity may not be detectable experimentally. Furthermore, the placement of the line scan in imaging experiments may not overlap precisely on the dyad and the resolution of the microscope which is ≈ 0.4 um is roughly similar to the size of a dyad (37,38).

## 5 Conclusions

By incorporating the behaviour of both RyRs and IP_3_Rs in our 1D model of the dyad, we show that IP_3_R activity elevates dyadic [Ca^2+^], which sensitizes proximal RyRs towards activation. The colocalization of IP_3_Rs with RyRs in the dyad thus increases the propensity for RyR-mediated Ca^2+^ sparks which potentially underlies the ECC-modulating effects seen in ventricular cardiomyocytes treated with G_q_ agonists. In this regard, further work is needed to link our findings of IP_3_R-influenced Ca^2+^ spark formation to multiscale whole-cell cardiomyocyte models incorporating IP_3_ signalling (39) and Ca^2+^ cycling (40,41) to elucidate its overall impact on global cytosolic Ca^2+^ transient dynamics and ECC (42,43).

## Acknowledgements

This research was supported in part by the Australian Government through the Australian Research Council Discovery Projects funding scheme (project DP170101358) to EJC and VR, the Australian Research Council Centre of Excellence in Convergent Bio-Nano Science and Technology (project CE140100036) to EJC, the KU Leuven Global PhD Partnerships with The University of Melbourne Grant (GPUM/21/036) to VR and HLR, and The University of Melbourne’s Research Computing Services and the Petascale Campus Initiative. HLR wishes to acknowledge financial support from the Research Foundation Flanders (FWO) through Project Grant G08861N and Odysseus programme Grant 90663. JC would like to thank Dr Pengxing Cao for the useful discussions on IP_3_R modelling. S.T.J. is supported by the Australian Research Council (Project No. DE200100988).

## Abbreviations

Ca^2+^: calcium
ECC: excitation contraction coupling
RyR: ryanodine receptor
IP_3_: inositol 1,4,5-trisphosphate
IP_3_R: IP_3_ receptor
IP_3_R-1: type 1 IP_3_R
IP_3_R-2: type 2 IP_3_R
LTCC: L-type Ca^2+^ channel
SR: sarcoplasmic reticulum
JSR: junctional SR
NSR: network SR
CICR: Ca^2+^-induced Ca^2+^ release
GPCR: G protein-coupled receptor
ET-1: endothelin-1
IICR: IP_3_-induced Ca^2+^ release
CaM: calmodulin
TnC: troponin C
CSQ: calsequestrin
SERCA: sarco/endoplasmic reticulum ATPase
1D: 1-dimensional
PSF: point spread function

## Reference

1. Wang SQ, Song LS, Lakatta EG, Cheng H. Ca2+ signalling between single L-type Ca2+ channels and ryanodine receptors in heart cells. Nature. 2001 Mar;410(6828):592–6.

2. Cheng H, Lederer WJ, Cannell MB. Calcium Sparks: Elementary Events Underlying Excitation-Contraction Coupling in Heart Muscle. Science. 1993;262(5134):740–4.

3. Bers DM. Cardiac excitation–contraction coupling. Nature. 2002 Jan;415(6868):198–205.

4. Lipp P, Laine M, Tovey SC, Burrell KM, Berridge MJ, Li W, et al. Functional InsP3 receptors that may modulate excitation–contraction coupling in the heart. Current Biology. 2000 Aug 1;10(15):939–S1.

5. Foskett JK, White C, Cheung KH, Mak DOD. Inositol Trisphosphate Receptor Ca2+ Release Channels. Physiological Reviews. 2007 Apr 1;87(2):593–658.

6. Berridge MJ. The Inositol Trisphosphate/Calcium Signaling Pathway in Health and Disease. Physiological Reviews. 2016 Oct 1;96(4):1261–96.

7. Signore S, Sorrentino A, Ferreira-Martins J, Kannappan R, Shafaie M, Del Ben F, et al. Inositol 1, 4, 5-Trisphosphate Receptors and Human Left Ventricular Myocytes. Circulation. 2013 Sep 17;128(12):1286–97.

8. Harzheim D, Movassagh M, Foo RSY, Ritter O, Tashfeen A, Conway SJ, et al. Increased InsP3Rs in the junctional sarcoplasmic reticulum augment Ca2+ transients and arrhythmias associated with cardiac hypertrophy. PNAS. 2009 Jul 7;106(27):11406–11.

9. Nakayama H, Bodi I, Maillet M, DeSantiago J, Domeier TL, Mikoshiba K, et al. The IP3 Receptor Regulates Cardiac Hypertrophy in Response to Select Stimuli. Circulation Research. 2010 Sep 3;107(5):659–66.

10. Proven A, Roderick HL, Conway SJ, Berridge MJ, Horton JK, Capper SJ, et al. Inositol 1,4,5-trisphosphate supports the arrhythmogenic action of endothelin-1 on ventricular cardiac myocytes. Journal of Cell Science. 2006 Aug 15;119(16):3363–75.

11. Domeier TL, Zima AV, Maxwell JT, Huke S, Mignery GA, Blatter LA. IP3 receptor-dependent Ca2+ release modulates excitation-contraction coupling in rabbit ventricular myocytes. American Journal of Physiology-Heart and Circulatory Physiology. 2008 Feb;294(2):H596–604.

12. Harzheim D, Talasila A, Movassagh M, Foo RSY, Figg N, Bootman MD, et al. Elevated InsP3R expression underlies enhanced calcium fluxes and spontaneous extra-systolic calcium release events in hypertrophic cardiac myocytes. Channels. 2010 Jan 1;4(1):67–71.

13. Moschella MC, Marks AR. Inositol 1,4,5-trisphosphate receptor expression in cardiac myocytes. Journal of Cell Biology. 1993 Mar 1;120(5):1137–46.

14. Gilbert G, Demydenko K, Dries E, Puertas RD, Jin X, Sipido K, et al. Calcium Signaling in Cardiomyocyte Function. Cold Spring Harb Perspect Biol. 2020 Mar 1;12(3):a035428.

15. Demydenko K, Sipido KR, Roderick HL. Ca2+ release via InsP3Rs enhances RyR recruitment during Ca2+ transients by increasing dyadic [Ca2+] in cardiomyocytes. Journal of Cell Science [Internet]. 2021 Jul 22 [cited 2021 Sep 22];134(14). Available from: https://doi.org/10.1242/jcs.258671

16. Wullschleger M, Blanch J, Egger M. Functional local crosstalk of inositol 1,4,5-trisphosphate receptor- and ryanodine receptor-dependent Ca2+ release in atrial cardiomyocytes. Cardiovascular Research. 2017 Apr 1;113(5):542–52.

17. Walker MA, Williams GSB, Kohl T, Lehnart SE, Jafri MS, Greenstein JL, et al. Superresolution Modeling of Calcium Release in the Heart. Biophysical Journal. 2014 Dec 16;107(12):3018–29.

18. Picht E, Zima AV, Shannon TR, Duncan AM, Blatter LA, Bers DM. Dynamic Calcium Movement Inside Cardiac Sarcoplasmic Reticulum During Release. Circulation Research. 2011 Apr 1;108(7):847–56.

19. Zima AV, Picht E, Bers DM, Blatter LA. Termination of Cardiac Ca2+ Sparks. Circulation Research. 2008 Oct 10;103(8):e105–15.

20. Williams GSB, Chikando AC, Tuan HTM, Sobie EA, Lederer WJ, Jafri MS. Dynamics of Calcium Sparks and Calcium Leak in the Heart. Biophysical Journal. 2011 Sep 21;101(6):1287–96.

21. Cannell MB, Kong CHT, Imtiaz MS, Laver DR. Control of Sarcoplasmic Reticulum Ca2+ Release by Stochastic RyR Gating within a 3D Model of the Cardiac Dyad and Importance of Induction Decay for CICR Termination. Biophysical Journal. 2013 May 21;104(10):2149–59.

22. Cao P, Tan X, Donovan G, Sanderson MJ, Sneyd J. A Deterministic Model Predicts the Properties of Stochastic Calcium Oscillations in Airway Smooth Muscle Cells. PLOS Computational Biology. 2014 Aug 14;10(8):e1003783.

23. Cao P, Donovan G, Falcke M, Sneyd J. A Stochastic Model of Calcium Puffs Based on Single-Channel Data. Biophysical Journal. 2013 Sep 3;105(5):1133–42.

24. Siekmann I, Wagner LE, Yule D, Crampin EJ, Sneyd J. A Kinetic Model for Type I and II IP3R Accounting for Mode Changes. Biophysical Journal. 2012 Aug 22;103(4):658–68.

25. Baddeley D, Jayasinghe ID, Lam L, Rossberger S, Cannell MB, Soeller C. Optical single-channel resolution imaging of the ryanodine receptor distribution in rat cardiac myocytes. PNAS. 2009 Dec 29;106(52):22275–80.

26. Shen X, van den Brink J, Hou Y, Colli D, Le C, Kolstad TR, et al. 3D dSTORM imaging reveals novel detail of ryanodine receptor localization in rat cardiac myocytes. The Journal of Physiology. 2019;597(2):399–418.

27. Kolstad TR, van den Brink J, MacQuaide N, Lunde PK, Frisk M, Aronsen JM, et al. Ryanodine receptor dispersion disrupts Ca2+ release in failing cardiac myocytes. Dietz HC, Vunjak-Novakovic G, editors. eLife. 2018 Oct 30;7:e39427.

28. Smith GD, Keizer JE, Stern MD, Lederer WJ, Cheng H. A Simple Numerical Model of Calcium Spark Formation and Detection in Cardiac Myocytes. Biophysical Journal. 1998 Jul 1;75(1):15–32.

29. Rüdiger S, Shuai JW, Huisinga W, Nagaiah C, Warnecke G, Parker I, et al. Hybrid Stochastic and Deterministic Simulations of Calcium Blips. Biophysical Journal. 2007 Sep 15;93(6):1847–57.

30. Laver DR, Kong CHT, Imtiaz MS, Cannell MB. Termination of calcium-induced calcium release by induction decay: An emergent property of stochastic channel gating and molecular scale architecture. Journal of Molecular and Cellular Cardiology. 2013 Jan 1;54:98–100.

31. Higazi DR, Fearnley CJ, Drawnel FM, Talasila A, Corps EM, Ritter O, et al. Endothelin-1-Stimulated InsP3-Induced Ca2+ Release Is a Nexus for Hypertrophic Signaling in Cardiac Myocytes. Molecular Cell. 2009 Feb 27;33(4):472–82.

32. Blanch i Salvador J, Egger M. Obstruction of ventricular Ca2+-dependent arrhythmogenicity by inositol 1,4,5-trisphosphate-triggered sarcoplasmic reticulum Ca2+ release. The Journal of Physiology. 2018;596(18):4323–40.

33. Pinali C, Malik N, Davenport JB, Allan LJ, Murfitt L, Iqbal MM, et al. Post-Myocardial Infarction T-tubules Form Enlarged Branched Structures With Dysregulation of Junctophilin-2 and Bridging Integrator 1 (BIN-1). Journal of the American Heart Association. 2017 May 4;6(5):e004834.

34. Guo A, Zhang C, Wei S, Chen B, Song LS. Emerging mechanisms of T-tubule remodelling in heart failure. Cardiovasc Res. 2013 May 1;98(2):204–15.

35. Groff JR, Smith GD. Calcium-dependent inactivation and the dynamics of calcium puffs and sparks. Journal of Theoretical Biology. 2008 Aug 7;253(3):483–99.

36. Macquaide N, Tuan HTM, Hotta J ichi, Sempels W, Lenaerts I, Holemans P, et al. Ryanodine receptor cluster fragmentation and redistribution in persistent atrial fibrillation enhance calcium release. Cardiovascular Research. 2015 Dec 1;108(3):387–98.

37. Rog-Zielinska EA, Moss R, Kaltenbacher W, Greiner J, Verkade P, Seemann G, et al. Nano-scale morphology of cardiomyocyte t-tubule/sarcoplasmic reticulum junctions revealed by ultra-rapid high-pressure freezing and electron tomography. Journal of Molecular and Cellular Cardiology. 2021 Apr 1;153:86–92.

38. Hayashi T, Martone ME, Yu Z, Thor A, Doi M, Holst MJ, et al. Three-dimensional electron microscopy reveals new details of membrane systems for Ca2+ signaling in the heart. Journal of Cell Science. 2009 Apr 1;122(7):1005–13.

39. Cooling M, Hunter P, Crampin EJ. Modeling Hypertrophic IP3 Transients in the Cardiac Myocyte. Biophysical Journal. 2007 Nov 15;93(10):3421–33.

40. Hunt H, Tilūnaitė A, Bass G, Soeller C, Roderick HL, Rajagopal V, et al. Ca2+ Release via IP3 Receptors Shapes the Cardiac Ca2+ Transient for Hypertrophic Signaling. Biophys J. 2020 Sep 15;119(6):1178–92.

41. Vierheller J, Neubert W, Falcke M, Gilbert S, Chamakuri N. A multiscale computational model of spatially resolved calcium cycling in cardiac myocytes: from detailed cleft dynamics to the whole cell concentration profiles. Frontiers in Physiology [Internet]. 2015 [cited 2022 Feb 15];6. Available from: https://www.frontiersin.org/article/10.3389/fphys.2015.00255

42. Crampin EJ, Smith NP, Hunter PJ. Multi-scale modelling and the IUPS physiome project. Histochem J. 2004 Sep 1;35(7):707–14.

43. Terkildsen JR, Niederer S, Crampin EJ, Hunter P, Smith NP. Using Physiome standards to couple cellular functions for rat cardiac excitation–contraction. Experimental Physiology. 2008;93(7):919–29.

44. Bers DM. Sources and Sinks of Activator Calcium. In: Bers DM, editor. Excitation-Contraction Coupling and Cardiac Contractile Force [Internet]. Dordrecht: Springer Netherlands; 2001 [cited 2022 Mar 16]. p. 39–62. Available from: https://doi.org/10.1007/978-94-010-0658-3_3

45. Wagner LE, Yule DI. Differential regulation of the InsP_3_ receptor type-1 and -2 single channel properties by InsP_3_, Ca^2+^ and ATP. J Physiol. 2012 Jul 15;590(14):3245–59.

46. Tran K, Smith NP, Loiselle DS, Crampin EJ. A Thermodynamic Model of the Cardiac Sarcoplasmic/Endoplasmic Ca2+ (SERCA) Pump. Biophysical Journal. 2009 Mar 4;96(5):2029–42.

