## Supplementary Text for "IP_3_R activity increases frequency of RyR-mediated sparks by elevating dyadic Ca^2+^"

### 7 Supplementary Materials

#### 7.1 Parameter Values

Values for every parameter used to simulate the $Ca^{2+}$ reaction diffusion in our model are shown in Table 1 and Table 2. All values are adapted directly from (17) except where otherwise indicated.

*Table 1. Parameter values of all species involved in reaction diffusion.*

| Species | Concentration (µM) | Diffusivity, $\mathcal{D}$ (µm$^2$/ms) | Forward Reaction Rate, $k_{on}$ (µM$^{-1}$ms$^{-1}$) | Backward Reaction Rate, $k_{off}$ (ms$^{-1}$) |
|---|---|---|---|---|
| $[Ca^{2+}]_c$ | 0.1 (initial) | 0.22 | - | - |
| $[Ca^{2+}]_{JSR}$ | 1000 (initial) | 0.35[1] | - | - |
| $[Ca^{2+}]_{NSR}$ | 1000 (initial) | 0.06 | - | - |
| ATP | 455 (total) | 0.14 | 0.225 | 45 |
| CaM | 24 (total) | 0.025 | 0.025 | 0.238 |
| Fluo-4 | 100 (total) | 0.042 | 0.0488[1] | 0.0439[1] |
| TnC | 70 (total) | 0 | 0.039 | 0.02 |
| CSQ | 30000 (total) | 0 | 0.1 | 63.8 |

*Table 2. Parameter values involved in calculating $Ca^{2+}$-handling protein fluxes and JSR refill.*

| $Ca^{2+}$-Handling Protein | Parameter | Description | Value |
|---|---|---|---|
| RyR | $g_{RyR}$ | RyR $Ca^{2+}$ release flux rate | 2.8 ms$^{-1}$ [2] |
| $IP_3R$ | $g_{IP_3R}$ | $IP_3R$ $Ca^{2+}$ release flux rate | 0.982 ms$^{-1}$ [3] |
| SERCA | $A_p$ | SERCA concentration | 75 µM[4] |
| | $K_{D_c}$ | SERCA sensitivity to $[Ca^{2+}]_c$ | 910 µM |
| | $K_{D_{SR}}$ | SERCA sensitivity to $[Ca^{2+}]_{NSR}$ | 2240 µM |
| JSR | $g_{refill}$ | JSR refill flux rate | 0.20 ms$^{-1}$ [5] |

#### 7.2 RyR Model

The RyR model used in our simulations is directly adapted from that developed by (21). The gating of each RyR is modelled as a 2-state Markov process (**Figure 4**).

$$CLOSE \underset{k_{close}}{\overset{k_{open}}{\rightleftharpoons}} OPEN$$

***Figure 4. State diagram of RyR model.*** *Developed by (21), this model of the RyR consists of 2 states, denoted by OPEN and CLOSE, that the RyR transitions between at transition rates $k_{open}$ and $k_{close}$.*

Where the $Ca^{2+}$-dependent transition rates, in ms, between the states, are expressed as,

---

[1] Value taken from (21)

[2] Adjusted to give a realistic $Ca^{2+}$ spark profile

[3] Value calculated as 2.85 times lower than $g_{RyR}$ as $Ca^{2+}$ conductance of $IP_3Rs$ is estimated to be ~2.85 lower than RyRs (5)

[4] Value taken from (44)

[5] Adjusted to give a $[Ca^{2+}]_{JSR}$ exponential recovery time constant of ~130 ms as in (17–19)

$$k_{open} = \min(3.17 \times 10^2 \times [Ca^{2+}]_c^{2.8}, 0.7)$$
$$k_{close} = \max(0.25 \times [Ca^{2+}]_c^{-0.5}, 0.9)$$

#### 7.3 $IP_3R$ Model

$IP_3Rs$ in our simulations are modelled after that developed by (23) who modified the park-drive model (24) to account for unsteady state kinetics of $IP_3Rs$ when subject to constantly changing concentrations of regulatory ligands (in this case, $Ca^{2+}$). The gating of each $IP_3R$ is modelled as a 6-state Markov process (**Figure 5**).

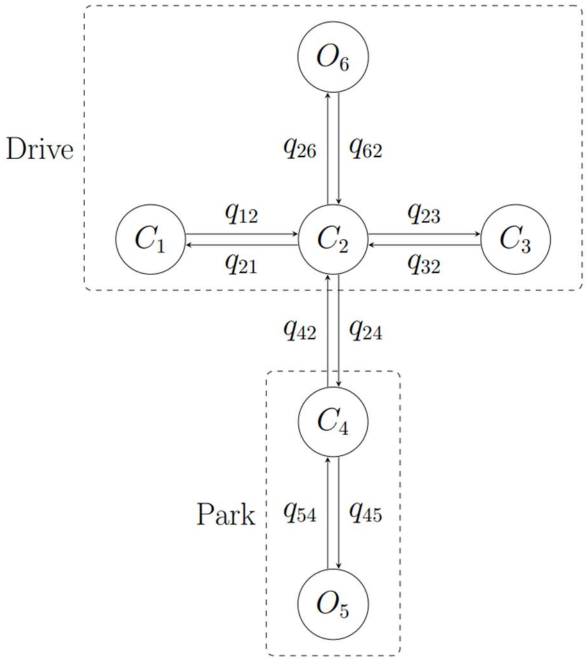


***Figure 5. State diagram of $IP_3R$ model.*** *Developed by (24), this model of the $IP_3R$ consists of six states that are categorised into two modes of activity: Park and Drive. Park mode is when the channel is at the closed state $C_4$ or open state $O_5$. Drive mode is when the channel is at closed states $C_1$, $C_2$, $C_3$ or open state $O_6$. State transition rates are denoted by $q$. Intramodal transition rates are constants whereas intermodal transition rates are dependent on ligand concentration.*

Intramodal transition rates are constants whose values are shown in Table 3 below,

*Table 3. Constant $IP_3R$-2 transition rates. Values obtained from (24).*

| $IP_3R$-2 State Transition Rate | Value ($ms^{-1}$) |
|---|---|
| $q_{12}$ | 1.14 |
| $q_{21}$ | 0.0958 |
| $q_{23}$ | 0.0047 |
| $q_{32}$ | 0.0119 |
| $q_{26}$ | 10.100 |
| $q_{62}$ | 3.270 |
| $q_{45}$ | 0.0041 |
| $q_{54}$ | 3.420 |

Intermodal transition rates $q_{24}$ and $q_{42}$ are ligand-dependent whose expressions are given by,

$$q_{24} = a_{24} + V_{24}(1 - m_{24}h_{24})$$
$$q_{42} = a_{42} + V_{42}m_{42}h_{42}$$

Where variables $a$, $V$, $m$, and $h$ are functions of ligand concentrations $IP_3$, $[IP_3]$ and $Ca^{2+}$, $[Ca^{2+}]$ and are given by the following expressions. These expressions take a similar form to that in (22,23).

$$a_{24} = \frac{100}{[IP_3]^{54.5} + 0.923^{54.5}}$$
$$a_{42} = 1.0 + \frac{24.5}{[IP_3]^{2.8} + 3.4^{2.8}}$$
$$V_{24} = 200.3 + \frac{24.1[IP_3]^{54.9}}{[IP_3]^{54.9} + 46.8^{54.9}}$$
$$V_{42} = 60.0 + \frac{745.0}{[IP_3]^{8.6} + 1.0^{8.6}}$$
$$m_{24} = \frac{[Ca^{2+}]^{n_{24}}}{k_{24}^{n_{24}} + [Ca^{2+}]^{n_{24}}}$$
$$m_{42} = \frac{[Ca^{2+}]^{n_{42}}}{k_{42}^{n_{42}} + [Ca^{2+}]^{n_{42}}}$$
$$h_{24} = \frac{k_{-2}^{n_{-24}}}{k_{-24}^{n_{-24}} + [Ca^{2+}]^{n_{24}}}$$
$$h_{42} = \frac{k_{-42}^{n_{-42}}}{k_{-42}^{n_{-42}} + [Ca^{2+}]^{n_{42}}}$$
$$k_{24} = 0.0358$$
$$k_{42} = 0.15$$
$$n_{24} = 9.5$$
$$n_{42} = 5.5$$
$$k_{-24} = 15.9 + \frac{774.2}{[IP_3]^{2.7} + 33.0^{2.7}}$$
$$k_{-42} = 0.8 + \frac{19000}{[IP_3]^{11.6} + 86.8^{11.6}}$$
$$n_{-24} = 1.14 + \frac{1.19[IP_3]^{1.25}}{[IP_3]^{1.25} + 20.7^{1.25}}$$
$$n_{-42} = 1.7 + \frac{37.8}{[IP_3]^{15.1} + 1.2^{15.1}}$$

Coefficients and exponents in expressions of the gating variables stated above are determined by fitting the curve of $q_{24}$ and $q_{42}$ to their known steady state data points. These data points (**Figure 6**A) were previously derived from experimental data by (24) and are specific to $IP_3R$-2. We chose to fit our plots of $q_{24}$ and $q_{42}$ to data points obtained at 1 µM and 10 µM [$IP_3$] and 5 mM [ATP] as there were more data points that we could fit our curves to and also because 5 mM [ATP] was closer to the physiological [ATP] in cardiomyocytes. The resultant fitted curves of $q_{24}$ and $q_{42}$ as a function of [$Ca^{2+}$] and [$IP_3$] are shown in **Figure 6**A. $q_{24}$ and $q_{42}$ at 0.15 µM [$IP_3$], the concentration at which $IP_3$ is fixed in all our simulations, were then extrapolated from these expressions and is shown in **Figure 6**B. The corresponding open probability curves calculated (**Figure 6**C) are comparable to those obtained from experiments (45).

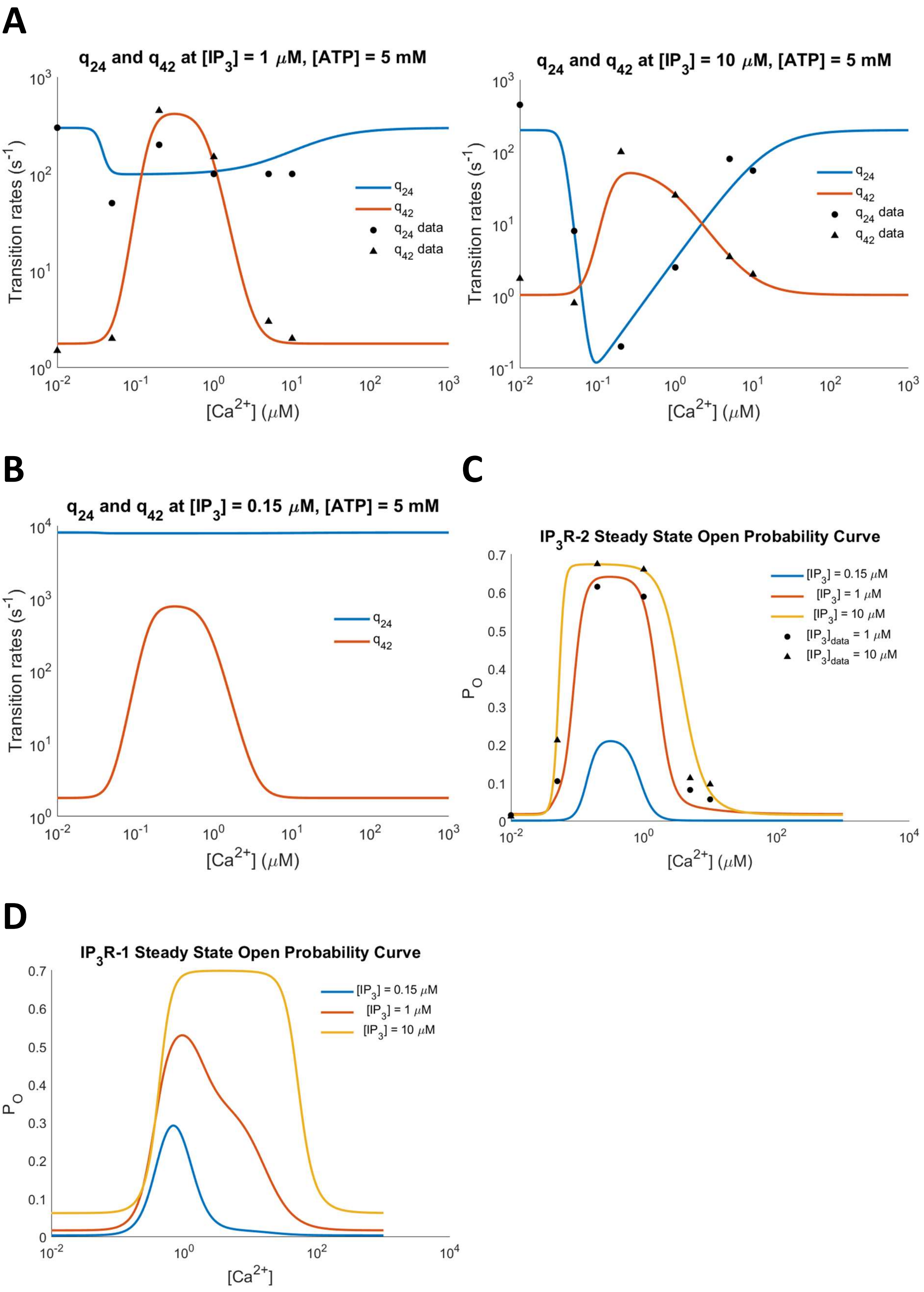


***Figure 6. Steady state and fitted plots of $q_{24}$ and $q_{42}$. A:*** *Plots were obtained by adjusting coefficients and exponents in expressions for variables $a$, $V$, $m$, $h$, and $k$ to fit steady state $q_{24}$ and $q_{42}$ data points obtained from (24).* ***B:*** *Steady state $q_{24}$ and $q_{42}$ plot at an $IP_3$ concentration of 0.15 μM.* ***C:*** *The corresponding open probability curve of $IP_3R$-2.* ***D:*** *The open probability curve of $IP_3R$-1 model developed in (22).*

To account for $IP_3R$-2 gating behaviour in an environment where [$Ca^{2+}$] is constantly changing, the non-steady state kinetics of the $Ca^{2+}$-dependent gating variables were assumed to obey the differential equation of the form below (23),

$$\frac{dG}{dt} = \lambda_G(G_\infty - G)$$

Where, $G$ represent the current value of gating variables $m_{24}$, $h_{24}$, $m_{42}$, and $h_{42}$ and $G_\infty$ represents the value of the same variables at steady state. $\lambda_G$ is the equilibrium approach rate whose values are given in Table 4.

*Table 4. The equilibrium approach rate for all $Ca^{2+}$-dependent gating variables. Values are obtained from (22,23).*

| Equilibrium Approach Rate | Value ($ms^{-1}$) |
|---|---|
| $\lambda_{m_{24}}$ | 0.1 |
| $\lambda_{h_{24}}$ | 0.04 |
| $\lambda_{m_{42}}$ | 0.1 |
| $\lambda_{h_{42}}$ | 0.1 when $IP_3R$ is open, $5 \times 10^{-4}$ when closed |

#### 7.4 SERCA Model

The SERCA model implemented and its parameters were directly adapted from (20) which based it on the simplified thermodynamically realistic model developed by (46). The $Ca^{2+}$ uptake flux by SERCA, $J_{SERCA}$, is given by

$$J_{SERCA} = 2v_{cycle}A_p$$

Where each term is defined as:

$$v_{cyc} = \frac{3.24873 \times 10^{12}K_c^2 + K_c(9.17846 \times 10^6 - 11478.2K_{SR}) - 0.329904K_{SR}}{D_{cycle}}$$

$$D_{cycle} = 0.104217 + 17.293K_{SR} + K_c(1.75583 \times 10^6 + 7.61673 \times 10^6 K_{SR}) + K_c^2(6.08462 \times 10^{11} + 4.50544 \times 10^{11}K_{SR})$$

$$K_c = \left(\frac{[Ca^{2+}]_c}{K_{D_c}}\right)^2$$

$$K_{SR} = \left(\frac{[Ca^{2+}]_{NSR}}{K_{D_{SR}}}\right)^2$$

$v_{cycle}$ corresponds to the cycling rate per SERCA molecule while $A_p$ corresponds to the cytosolic concentration of SERCA molecules. $K_{D_c}$ and $K_{D_{SR}}$ are constants quantifying the sensitivity of SERCA activity to $[Ca^{2+}]_c$ and $[Ca^{2+}]_{NSR}$ respectively. Their values are given in Table 2.

#### 7.5 $Ca^{2+}$ Spark Analysis

$Ca^{2+}$ releases at the dyad are identified as $Ca^{2+}$ sparks when it involves the opening of ≥ 7 RyRs in the dyad. This classification is justified as $Ca^{2+}$ sparks that occur in our simulations typically involve the opening of 12 – 20 RyRs in the dyad. The amplitude and FDHM of $Ca^{2+}$ sparks were then obtained from $Ca^{2+}$ trace of $Ca^{2+}$ spark events using the *findpeaks* function in MATLAB (**Figure** 7B, C).

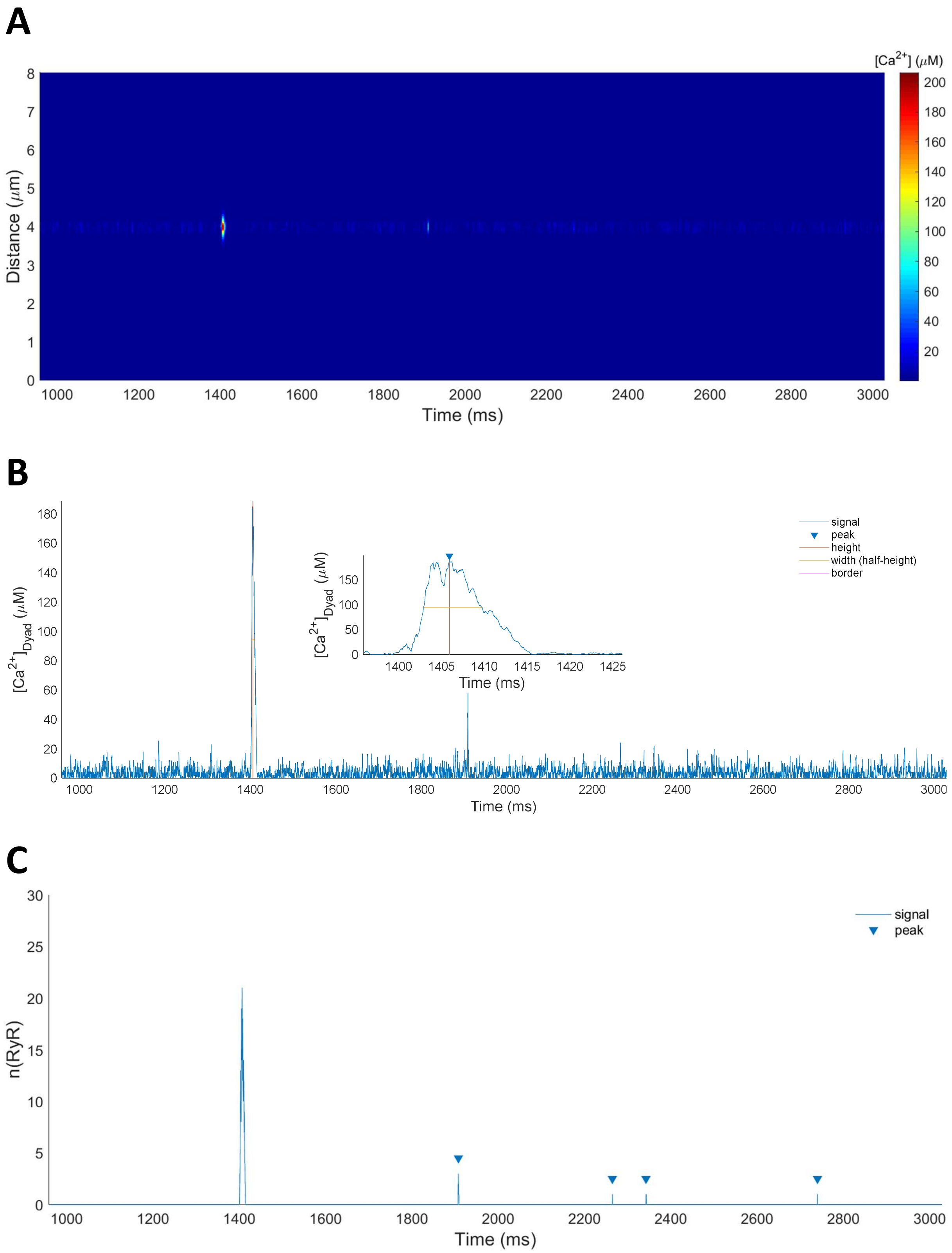


***Figure 7. Detection and analysis of $Ca^{2+}$ spark events. A:*** *The [$Ca^{2+}$] equivalent of a line scan image.* ***B:*** *$Ca^{2+}$ Spark detection and measurement of its amplitude and FDHM. $Ca^{2+}$ spark detected are denoted by an inverted triangle at its peak [$Ca^{2+}$]. Inset shows how the amplitude and FDHM of a detected $Ca^{2+}$ spark is measured.* ***C:*** *Detection of spontaneous RyR openings that do not develop into a full $Ca^{2+}$ spark. Notice that full $Ca^{2+}$ spark events were excluded from detection.*

#### 7.6 $Ca^{2+}$ Spark Fluorescence

**Figure 8**A shows the simulated fluorescence line scan images and traces from the center of the dyad together with their equivalent [$Ca^{2+}$] counterpart **Figure 8**B. Due to the 1D nature of our model, we convolved the simulated fluorescence with a 1D Gaussian PSF with a FWHM of 0.41 μm. Notice the plateau in the fluorescence trace of a $Ca^{2+}$ spark that indicates the saturation of the indicator dye. Our simulated fluorescence result shows a similar $Ca^{2+}$ spark amplitude independent of $IP_3R$ activity, which is consistent with experimental data (15). However, we are unable to reliably conclude this as it may be biased by the saturation of the indicator dye.

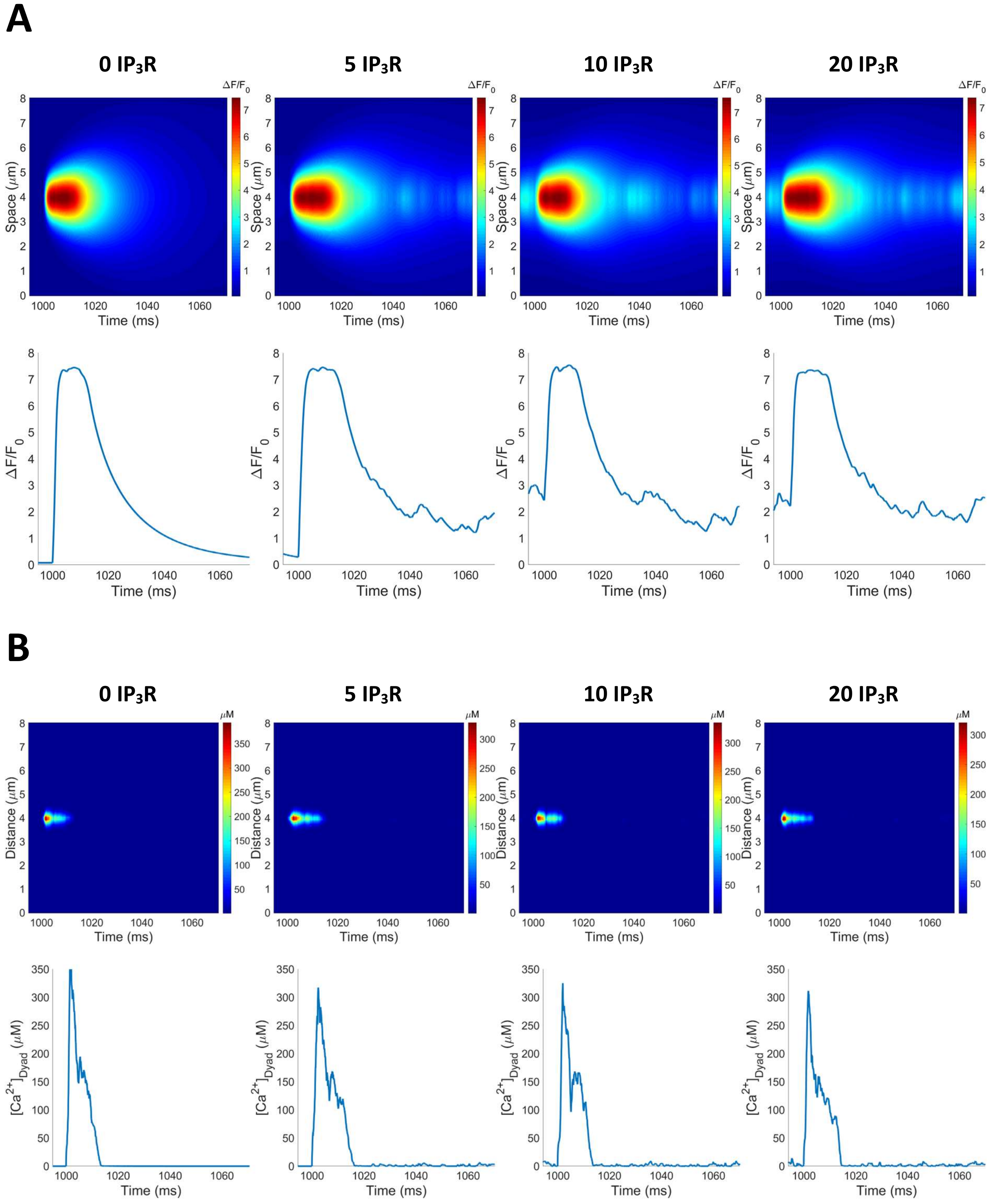

***Figure 8. $Ca^{2+}$ spark fluorescence and its underlying [$Ca^{2+}$]. A:*** *Line scan images of a $Ca^{2+}$ spark and its fluorescence trace taken at the center of the line scan.* ***B:*** *The [$Ca^{2+}$] equivalent of a line scan image and its [$Ca^{2+}$] trace taken at the center of the line scan.*
